## Supplementary material for "Atypical integrative element with strand-biased circularization activity assists interspecies antimicrobial resistance gene transfer from *Vibrio alfacsensis*": S1_Table: S1_Table.docx

| Strain | Copy number of SE-6945 | MIC (ug/ ml) | Genotype |
| --- | --- | --- | --- |
| LN27 | 3 | 1000 | *insJ*::SE-6945::pSEA2 *yjjNt*::SE-6945 |
| LN58 | 2 | 500 | *insJ*::SE-6945::pSEA2 |
| LN84 | 2 | 500 | *insJ*::SE-6945::pSEA2 |
| LN89 | 2 | 500 | *insJ*::SE-6945::pSEA2 |
| LN91 | 2 | 500 | *yjjNt*::SE-6945::pSEA2 |
| LN33 | 2 | 500 | *insJ*:: SE-6945::pSEA2 |
| LN34 | 2 | 500 | *insJ*:: SE-6945::pSEA2 |
| LN35 | 2 | 500 | *insJ*::SE-6945::pSEA2 |
| LN28 | 1 | 500 | chr::pSEA2 |
| LN29 | 2 | 250 | *yjjNt*::SE-6945::pSEA2 |
| LN30 | 2 | 250 | *yjjNt*::SE-6945::pSEA2 |
| LN82 | 2 | 250 | *yjjNt*::SE-6945::pSEA2 |
| LN88 | 2 | 250 | *yjjNt*::SE-6945::pSEA2 |
| LN52 | 1 | 125 | *yjjNt*::SE-6945 |
| LN86 | 1 | 62.5 | chr::pSEA2 |
| LN32 | 1 | 62.5 | chr::pSEA2 |
| LN36 | 1 | 16 | chr::pSEA2 |
| LN90 | 2 | 16 | *insJ*::SE-6945::pSEA2 |
| JW0452 | 0 | 2 | Parent |
