## Supplementary material for "Atypical integrative element with strand-biased circularization activity assists interspecies antimicrobial resistance gene transfer from *Vibrio alfacsensis*": S2_Table: S2_Table.docx

| Organisms | Data type | GenBank/SRA/DRA accession numbers or figshare link |
| --- | --- | --- |
| *V. alfacsensis* 04Ya249 | Pacbio SRII reads | DRA008632 |
| *V. alfacsensis* 04Ya108 | Pacbio SRII reads | DRA011098 |
| *E. coli* TJ249 | Pacbio SRII reads | DRA013462 |
| *V. alfacsensis* LN95 | Illumina raw reads | DRA011762 |
| *V. alfacsensis* 04Ya249 | Complete sequence of Chromosome 1 | AP019849 |
| *V. alfacsensis* 04Ya249 | Complete sequence of Chromosome 2 | AP019850 |
| *V. alfacsensis* 04Ya249 | Complete sequence of pVA249 | AP019852 |
| *V. alfacsensis* 04Ya249 | Complete sequence of pSEA2 | AP019851 |
| *V. alfacsensis* 04Ya108 | Complete sequence of Chromosome 1 | AP024165 |
| *V. alfacsensis* 04Ya108 | Complete sequence of Chromosome 2 | AP024166 |
| *V. alfacsensis* 04Ya108 | Complete sequence of pYa108 | AP024168 |
| *V. alfacsensis* 04Ya108 | Complete sequence of pSEA1 | AP024167 |
| *E. coli* TJ249 | Assembly | https://doiorg/106084/m9figshare13332467 |
