## Supplementary figures and images for "Atypical integrative element with strand-biased circularization activity assists interspecies antimicrobial resistance gene transfer from *Vibrio alfacsensis*"

### S1_FIG.tif

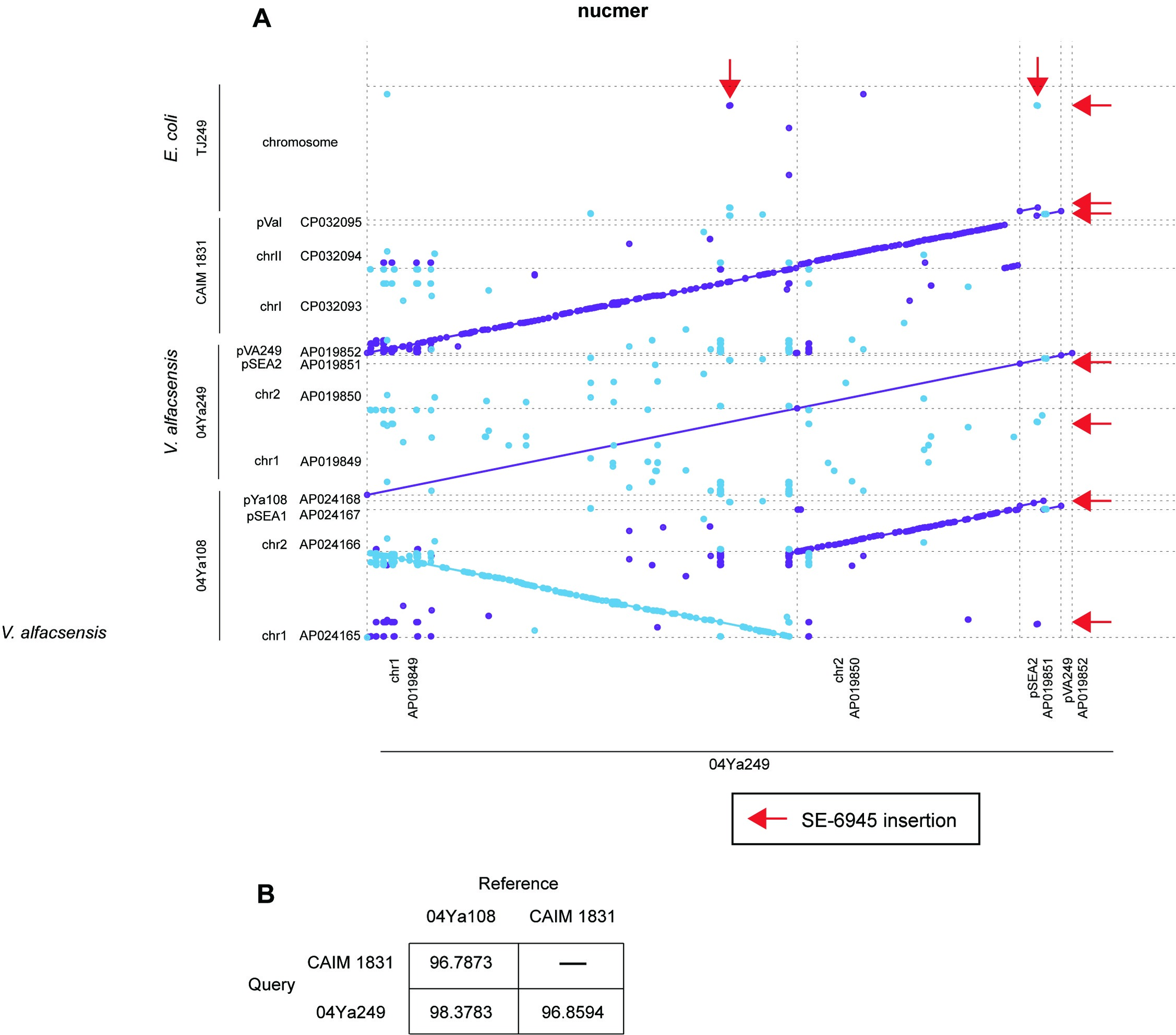

### S2_FIG.tif

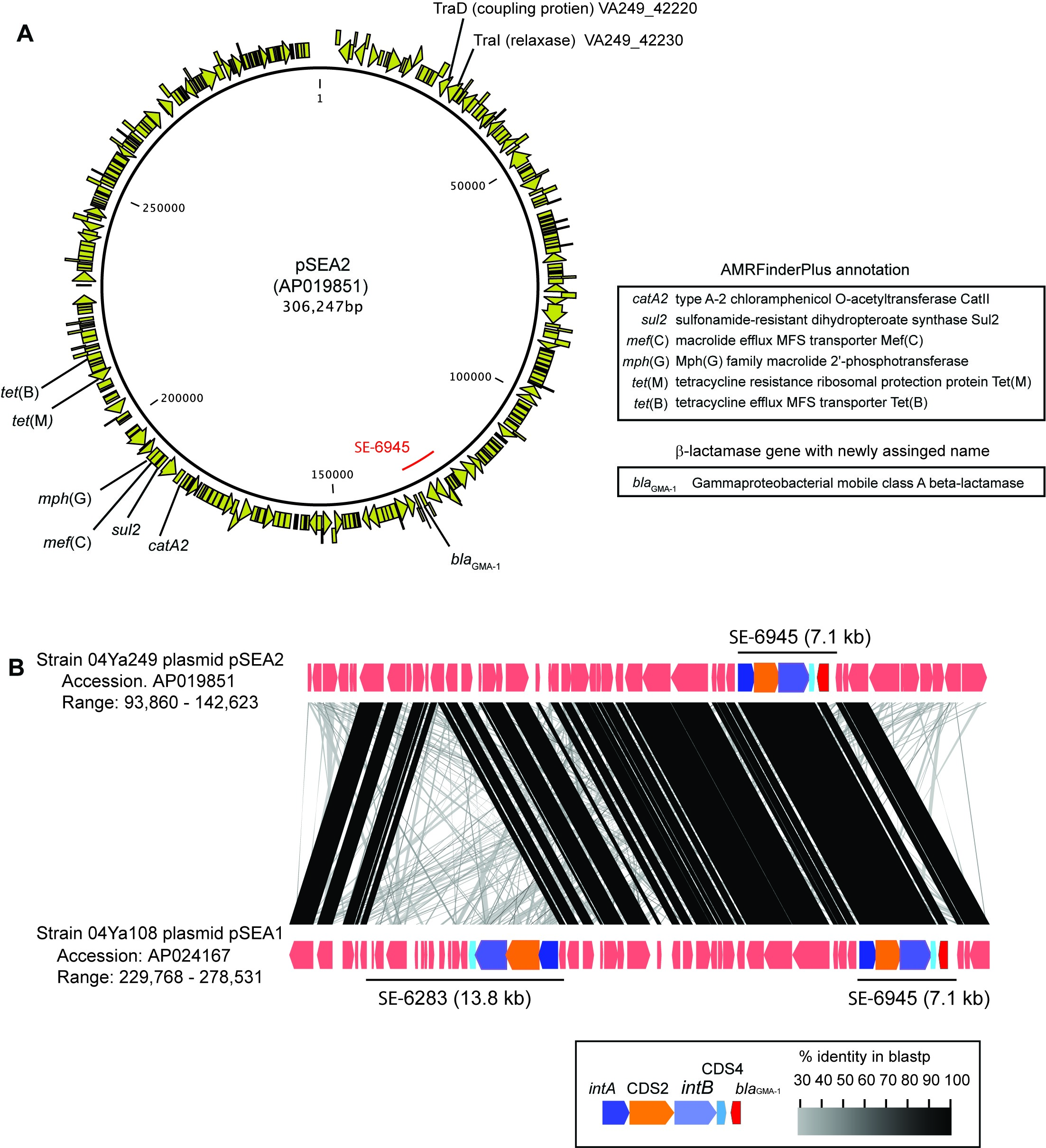

### S5_FIG.tif

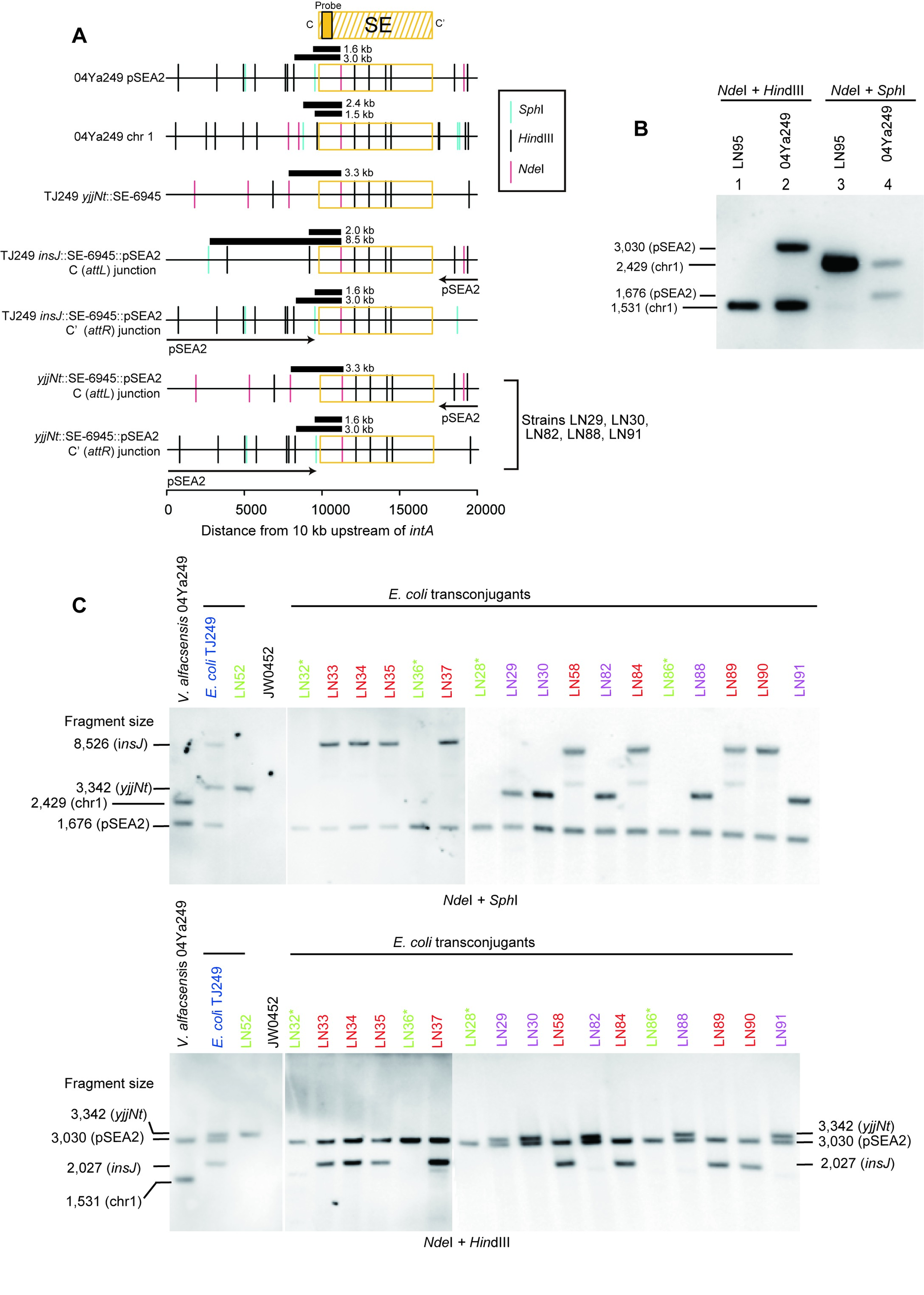

### S8_FIG.tif

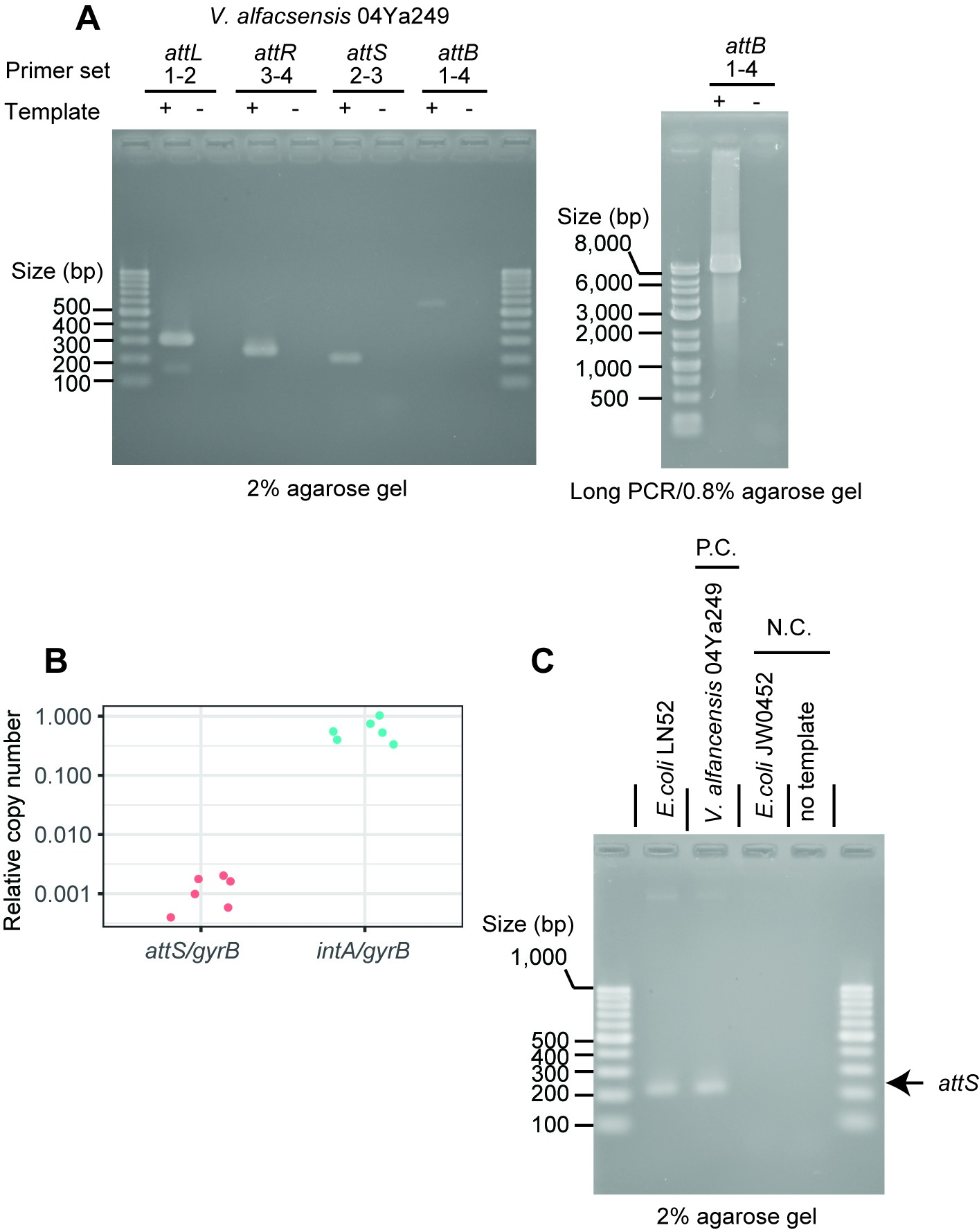
