## Supplementary material for "Atypical integrative element with strand-biased circularization activity assists interspecies antimicrobial resistance gene transfer from *Vibrio alfacsensis*": S3_FIG: S3_FIG.docx

Putative *yjjNt* transcriptional terminator

```
                                1234567890123456789012345678901234567890123456789012345678901234567890
attL_SE-6945_pSEA2_AP019851.1:  TAAAGCCTGATAAAGAAACCCGACAAGTGAACCATCTAATGCAACCCATGGTGCGCCTTTTCATATTGGC 70
attL_SE-6945_Eco(insJ):         CTTTCAAGATCCTCAATGCGTCGGTCTTTTGACAGCTCCAATGCTGATGCCGCTTTTTCTGGATCAACTG 70
attL_SE-6945_Eco(yjjNt):        TTAATTAAAGGCGTAATTACTTTCTGATAAGGCGAGATTATTAAAGTTGCCATGCAGCGTCCGGGGAAGT 70
attL_SE-6945_Val_AP019849.1:    GAAAAAAGAAGATTAAGTTGATGCCAACCCATCGTTGAAAACTATCAACACGATTCAAAAAGCCATGCAG 70
attL_SE-VhaWXL538_CP045070.1:   AAGAAGACTAAGTGATGTTCACTGCCAATCATTGTTGAAAGCTATCAACACGATTCAAAAAGCCATGCAG 70
```

motif C

```
                                1234567890123456789012345678901234567890123456789012345678901234567890
attL_SE-6945_pSEA2_AP019851.1:  ATATCGCCATGTTTTTCTTACAGCCAACATATCTAAATAGTTGTTTGACATAGATAACAGTATCTGTGTT 140
attL_SE-6945_Eco(insJ):         ATATTGCAATGTTTCTTTTACAGCCAACATATCTAAATAGTTGTTTGACATAGATAACAGTATCTGTGTT 140
attL_SE-6945_Eco(yjjNt):        GTTGGGCGCTGTTTTTTTTACAGCCAACATATCTAAATAGTTGTTTGACATAGATAACAGTATCTGTGTT 140
attL_SE-6945_Val_AP019849.1:    ACAATGCATGGCTTTTTTTACAGCCAACATATCTAAATAGTTGTTTGACATAGATAACAGTATCTGTGTT 140
attL_SE-VhaWXL538_CP045070.1:   ACAATGCATGGCTTTTTTTACAGCCAACATATCTAAATAGTTGTTTGACATAGATAATAGTATCTATGTT 140
                                     **   * ** * **************************************** ******* ****
```

Putative start codon for *intA*

```
                                12345678901234567890123456789012345678901234567890123456789012345
attL_SE-6945_pSEA2_AP019851.1:  TTTATTTGATTTGTTTAGATTGATAGTCTAACTTTAATTTAGATAAGTTAATTGGGTATTGTATG 205
attL_SE-6945_Eco(insJ):         TTTATTTGATTTGTTTAGATTGATAGTCTAACTTTAATTTAGATAAGTTAATTGGGTATTGTATG 205
attL_SE-6945_Eco(yjjNt):        TTTATTTGATTTGTTTAGATTGATAGTCTAACTTTAATTTAGATAAGTTAATTGGGTATTGTATG 205
attL_SE-6945_Val_AP019849.1:    TTTATTTGATTTGTTTAGATTGATAGTCTAACTTTAATTTAGATAAGTTAATTGGGTATTGTATG 205
attL_SE-VhaWXL538_CP045070.1:   TTTATTTGATTAAATTAGATTGATAGTCTAACTTTAATTTAGATGAGTGGAATAGGCATTGTATG 205
                                ************  ****************************** ***  * * ** ********
```
