## Supplementary material for "Atypical integrative element with strand-biased circularization activity assists interspecies antimicrobial resistance gene transfer from *Vibrio alfacsensis*": S4_FIG: S4_FIG.docx

motif C’

```
                                1234567890123456789012345678901234567890123456789012345678901234567890
attR_SE-6945_pSEA2_AP019851.1:  TATTTTGTGTGTAGCCCTTGTGCGTAAAGGGATTCCTAACTTTTTTATCTAACTTTATGTTAAGGGTATT 70
attR_SE-6945_Eco(insJ):         TATTTTGTGTGTAGCCCTTGTGCGTAAAGGGATTCCTAACTTTTTTATCTAACTTTATGTTAAGGGTATT 70
attR_SE-6945_Eco(yjjNt):        TATTTTGTGTGTAGCCCTTGTGCGTAAAGGGATTCCTAACTTTTTTATCTAACTTTATGTTAAGGGTATT 70
attR_SE-6945_Val_AP019849.1:    TATTTTGTGTGTAGCCCTTGTGCGTAAAGGGATTCCTAACTTTTTTATCTAACTTTATGTTAAGGGTATT 70
attR_SE-VhaWXL538_CP045070.1:   TATTTTTTGTGTAGCCCTTGTGCGTAAAGGGATTCCTAACTTTTTTATCTAACTTTATGTTAAGGGTATT 70
                                ****** ***************************************************************


                                1234567890123456789012345678901234567890123456789012345678901234567890
attR_SE-6945_pSEA2_AP019851.1:  TTTTTGTTTTCGATGTCACTATTGAGCTTACGAAGAACATCAACATAAGCGGTACCGAGTGAATCGAATG 140
attR_SE-6945_Eco(insJ):         TTCTTGGTGCCAATCTTGAGCGCGCGTAAACCAGCTTCTCCGCGCTCTTCATAGACCTTCAGCCACCTGG 140
attR_SE-6945_Eco(yjjNt):        TTCTTGTTTCTTAATAATGTGTTGTAAGCCGTAGAAGGCGTGTAGGTCGCACCCTATGCGACCTACACAT 140
attR_SE-6945_Val_AP019849.1:    TTCTTGTACTCTAATGAAAAATAGAATTAGGGAAGTTTACAGCCATGCTTATCGTCGTTTCTCCAGCAAA 140
attR_SE-VhaWXL538_CP045070.1:   TTTTTGTACTCTAAATAGAAAATAGAATTAGGGAAGTTAAAGTCATGCTCATCGTAGTTTCTCCAGCTAA 140
                                ** ***      *


                                1234567890
attR_SE-6945_pSEA2_AP019851.1:  TACACTGGTT 150
attR_SE-6945_Eco(insJ):         CTACAGAACC 150
attR_SE-6945_Eco(yjjNt):        CAGTCAGGAA 150
attR_SE-6945_Val_AP019849.1:    GACACTTGAT 150
attR_SE-VhaWXL538_CP045070.1:   AACGCTCGAT 150
```
