## Supplementary material for "Atypical integrative element with strand-biased circularization activity assists interspecies antimicrobial resistance gene transfer from *Vibrio alfacsensis*": S6_FIG: S6_FIG.docx

motif C

```
                              1234567890123456789012345678901234567890123456789012345678901234567890
attL_SE-VroAM7_AP019798.1:    CGAAGACCATGCAGCTGCTGACCAGTTCGGCGTGTGGGGCGAGAAAAAGTTTACCCTTAGAAATTTTTTA 70
attL_SE-VscVS-12_CP016307.1:  CGAAGATCATGCCGTGGCAGAACAGTTTGGCGTTTGGGGCGAGAAGAAATTTACCCTTAGAAATTTTTCA 70
attL_SE-ValK09K1_CP017919.1:  TGAAGACCATGCCGTTGCTGACCAGTTCGGCGTTTGGGGCGAAAAGAAATTTACCCTTGATTATTTTCCA 70
attL_SE-6283_AP024165.1:      TGAAGATCATGCTGTTGCAGAACAGTTTGGTGTTTGGGGTGAAAAGAAGTTCACCCTTAGAAATTTTTTA 70
attL_SE-6283_Ecol_TJ108W0:    TGAGGACCACCAGGTGTGCGAACAATTCGGCGTCTGGGGTGAAAAGTCCTTCACCCTTAGAAATTTTTTA 70
attL_SE-6283_pSEA1_AP024167.1:TATCTCAAACGATGGCGTCTATGCTATTGATTCTCTTTATGAAGAAGAGGGCACCCTTAGAAATTTTTTA 70
                                      *    *      *     * *           **  *       ******    *****  *


                              1234567890123456789012345678901234567890123456789012345678901234567890
attL_SE-VroAM7_AP019798.1:    TGACAAAAAACCTTGATGTATTTTTATGACAAACCTATCTTTGATATGTGGGTAACATATCGAGGATTGT 140
attL_SE-VsCVS-12_CP016307.1:  TGACAAAAAACCTTGACGTTTTTTTATGACAATTCTATCTTTGATATGTGGGTAGCATATCGAGGATTGT 140
attL_SE-VAlK09K1_CP017919.1:  TGACAGCAAACCTTGATTAATTTTTATGACAAAACTACCTTTAATATGTGGGTAACATATTGAGGATTGT 140
attL_SE-6283_AP024165.1:      TGACATAAAACCTTGGATTTTTTTTATGACAAAACTATCTTTGATATGTGGCTAACATATCGAGGGTTGT 140
attL_SE-6283_Ecol_TJ108W0:    TGACATAAAACCTTGGATTTTTTTTATGACAAAACTATCTTTGATATGTGGCTAACATATCGAGGGTTGT 140
attL_SE-6283_pSEA1_AP024167.1:TGACATAAAACCTTGGATTTTTTTTATGACAAAACTATCTTTGATATGTGGCTAACATATCGAGGGTTGT 140
                              *****  ********     ************  *** **** ******** ** ***** **** ****
```

Putative start codon for *intA*

```
                              123456
attL_SE-VroAM7_AP019798.1:    TTTGTG 146
attL_SE-VsCVS-12_CP016307.1:  TTTGTG 146
attL_SE-VAlK09K1_CP017919.1:  TTTGTG 146
attL_SE-6283_AP024165.1:      CTTGTG 146
attL_SE-6283_Ecol_TJ108W0:    CTTGTG 146
attL_SE-6283_pSEA1_AP024167.1:CTTGTG 146
                               *****
```
