## Supplementary material for "Atypical integrative element with strand-biased circularization activity assists interspecies antimicrobial resistance gene transfer from *Vibrio alfacsensis*": S7_FIG: S7_FIG.docx

motif C’

```
                               1234567890123456789012345678901234567890123456789012345678901234567890
attR_SE-VroAM7_AP019798.1:     AACTCACTGTTAAAGATATAGTTTGTTATAAATCGTCAAAAATTGAGTCTTAAAAAATCCACAAGGGTAT 70
attR_SE-VscVS-12_CP016307.1:   AACTCACTGTTAAAGTTAGAGTTTGTTATAAATCGTCAGCAATTGAGTCTAAAAAAATTCCCAAGGGTAG 70
attR_SE-ValK09K1_CP017919.1:   AAACCACTATTAAAGAAAGATTTTTCATAAAACTGTCAAAGTTGTCGTCTTAGATTACTTCCAAGGGTGT 70
attR_SE-6283_AP024165.1:       AAAATACTGTTAGAGATAGATTTTTCTGCAAATTGTCATAAATTTAGTCTTAAAGAATTTCCAAGGGTAA 70
attR_SE-6283_Ecol_TJ108W0:     AAAATACTGTTAGAGATAGATTTTTCTGCAAATTGTCATAAATTTAGTCTTAAAGAATTTCCAAGGGTAA 70
attR_SE-6283_pSEA1_AP024167.1: AAAATACTGTTAGAGATAGATTTTTCTGCAAATTGTCATAAATTTAGTCTTAAAGAATTTCCAAGGGTAG 70
                               ***  *** *** **  * * ***     ***  ****    *   **** * *  *    *******


                               12345678901234567890123456789012345678901234567890123456789
attR_SE-VroAM7_AP019798.1:     TTTGTCATGGGTAAAGTCTACGATGGTCTTCACCGCATTAGCTTCCTAATCAACGAAGA 129
attR_SE-VsCVS-12_CP016307.1:   TTCATTATGGGTAAAGTGTATGATGGCTTGCACCGTATTAGCTTCCTAATCAACGAGCA 129
attR_SE-ValK09K1_CP017919.1:   AAGTACATGGGCAAAGTATACGACGGTCTTCACCGCATTAGCTTTCTGATCAATGAAGA 129
attR_SE-6283_AP024165.1:       GAGGGCATGGGTAAAGTGTACGACGGACTTCACCGCATCAGTTTCCTGATCAACGAGGA 129
attR_SE-6283_ECol_TJ108W0:     GAGGGCATGGGCAAAACCTACGATGGCATTCATCGCATCAGCTTCCTGATTGACGCTGA 129
attR_SE-6283_pSEA1_AP024167.1: AAGTTCAAAGGCGTCAAATATGAGTTCGTATACAACGATACTTTCCTCTTTAGCGCTGA 129
                                        **       ** **     *  *          ** **  *    *  **
```
